# Protocol for designing and interpreting minigene assays to validate candidate splice altering variants

**DOI:** 10.64898/2026.05.05.723105

**Authors:** Whitney Whitford, Suzanne M. Musgrave, Russell G. Snell, Jessie C. Jacobsen

**Affiliations:** School of Biological Sciences, Waipapa Taumata Rau | The University of Auckland, New Zealand; Centre for Brain Research, Waipapa Taumata Rau | The University of Auckland, New Zealand

## Abstract

Variants affecting RNA splicing are a major contributor to human disease, yet the consequences of variants outside of the canonical splice motifs are often difficult to determine. Here, we present a protocol for minigene-based evaluation of candidate splice-altering variants. The methodology described includes locus-specific insert design, commercial gene fragment synthesis, and long-read sequencing. The combined approach enables rapid assay development and nucleotide level resolution of the effect on splice isoforms *in vitro*, providing a scalable framework for functional validation of predicted cryptic splice variants.

**Graphical abstract:** 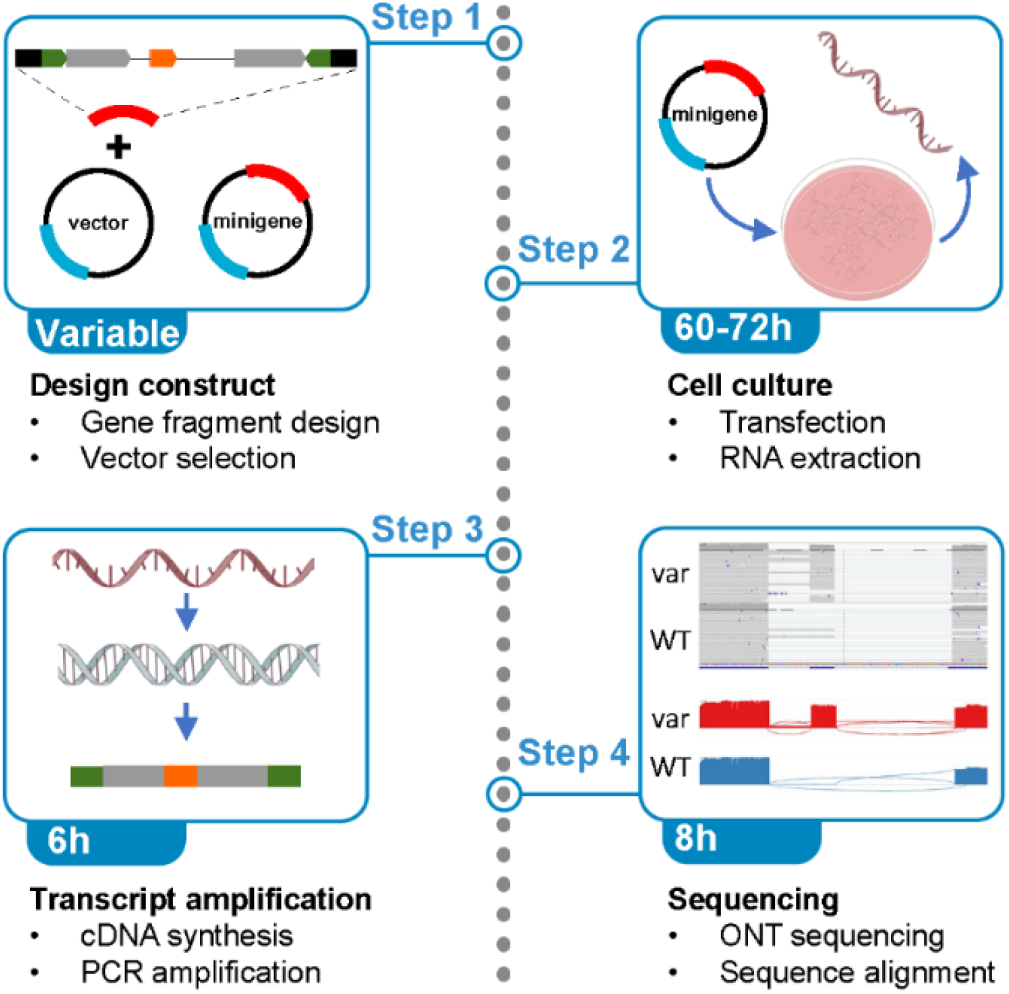

## Before you begin

### Background

Human genes are typically composed of short exons (average 309 bp) separated by longer introns (average 6,355 bp) that must be removed to process pre-mRNA into mature mRNA for translation[1], [2]. Accurate intron removal by the spliceosome requires recognition of conserved splice signals and regulatory elements at exon–intron junctions. Pre-mRNA splicing occurs through two sequential transesterification reactions. First, cleavage at the 5′ canonical splice site (beginning with a “GU” motif in ~95% of introns) is initiated by nucleophilic attack from an adenosine at the branch point (BP), located upstream of the 3′ splice site, forming a lariat intermediate. Downstream of the BP lies the polypyrimidine tract, which, together with the BP, helps define the 3′ canonical splice site (typically ending in a “AG” motif). In the second step, the upstream exon is ligated to the downstream exon, with the intron removed (Figure 1A).

**Figure 1.**
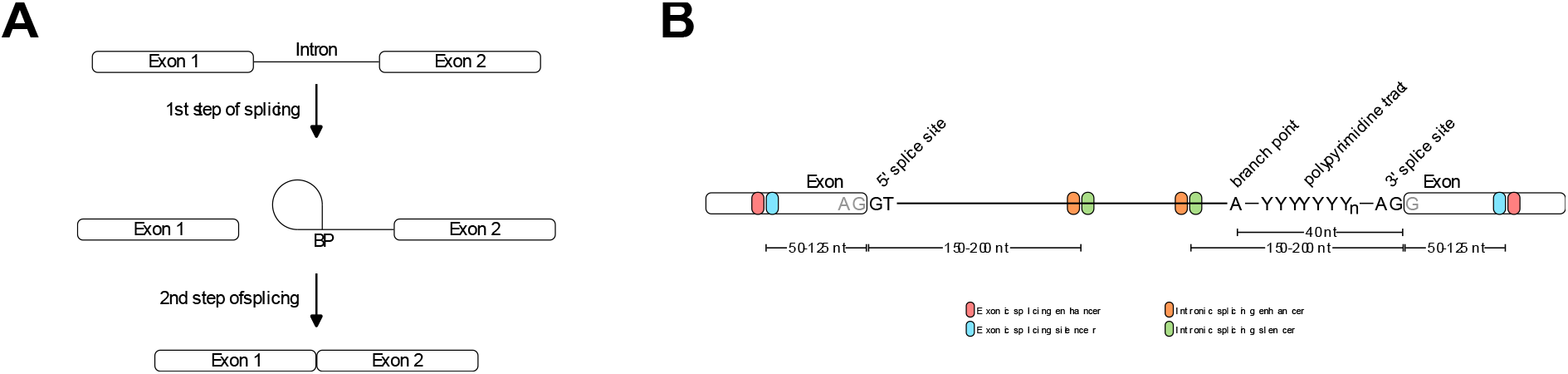
Splicing of nascent transcripts. A. Basic mechanism of splicing. Step 1 involves the cleavage of the 5’ canonical splice site with nucleophilic attack by the branch point. Step 2 involves ligation of the two exons together. B. Splicing elements and typical genomic locations. The key elements in splicing include the canonical 5’ and 3’ splice sites along with the intronic BP and polypyrimidine tract. Cis-acting enhancer and silencing elements are also present in exons and introns to regulate splicing.

In addition to these core splice signals (the 5′ canonical splice site, branch point sequence, polypyrimidine tract, and 3′ canonical splice site), splicing is further regulated by cis-acting elements, including exonic and intronic splicing enhancers (ESEs and ISEs) and silencers (ESSs and ISSs) (Figure 1B). These elements influence splice site selection of the spliceosome complex through interactions with trans-acting RNA-binding proteins. Variants that disrupt either core splice signals or regulatory elements can alter splice site recognition, resulting in aberrant transcript isoforms [3].

Variants affecting canonical splice donor (5′) or acceptor (3′) sites can lead to intron retention or exon skipping, respectively. Additionally, variants can create novel splice sites or strengthen existing non-canonical splice sites within intronic or exonic sequence, for example by increasing similarity to consensus splice site motifs and enhancing spliceosome recognition. This can result in activation of cryptic splice sites and, depending on sequence context, may lead to partial exon truncation or extension, or inclusion of a cryptic exon. In addition, genetic variants located within exonic or intronic splicing enhancers and silencers can disrupt normal splice site selection, often producing partial exon inclusion or exclusion [4], [5]. Variants affecting the branch point or polypyrimidine tract may also alter spliceosome splice site recognition, however, whether they have an effect, and the nature of any resulting splicing changes, are often difficult to predict [6].

Genetic variants affecting splicing are a major contributor to human disease. Approximately 15% of known disease-causing variants occur at canonical splice sites [3], [7], and this proportion may increase to as much as 60% when considering variants that disrupt splicing regulatory elements [4], [8]. Pathogenic variants affecting gene function through splicing effects have been identified and functionally validated in all types of core cis-acting elements, including exonic and intronic enhancers and silencers [6].

Genome-wide sequencing in clinical diagnostics, has dramatically expanded the mutation search space resulting a growing reliance on computational tools to predict the impact of candidate cryptic splice-altering variants [9], [10]. Although these tools, particularly those based on machine learning, demonstrate high sensitivity and specificity in predicting known splice-effecting variants, they may misclassify the presence, magnitude, or nature of splicing changes[9], and experimental validation is often required to confirm predicted pathogenic splicing effects in an appropriate biological context.

Plasmid transfection minigene assays provide an effective tool to evaluate the impact of genetic variants on splicing. These assays involve introducing a genomic fragment containing the variant of interest and flanking exonic and intronic sequence into a plasmid expression vector. This vector is then transiently transfected into cultured cells followed by RNA isolation, reverse transcription cDNA synthesis, PCR, and size-based analysis using gel electrophoresis or sequencing-based analysis revealing splicing outcomes. Traditionally, this required cloning of genomic fragments into minigene vectors, propagation in E. coli, and subsequent transfection[11]. However, advances in gene synthesis technologies now enable cost-effective, rapid and precise synthesis of minigene constructs by service providers, facilitating higher-throughput functional assessment of predicted splice variants.

Where available, RNA derived from patient cells provides the most physiologically relevant assessment of a patient-specific cryptic splice variant. However, access to disease-appropriate tissue is often limited, particularly for individuals with neurodevelopmental conditions or for genes that are not expressed in accessible tissues such as blood or fibroblasts. In these cases, minigene assays provide a practical and controlled alternative for functional validation.

While full-length gene reporter systems preserve broader endogenous splicing context[12], most human genes are prohibitively large for efficient synthesis, cloning, and transient transfection into mammalian cells[1], [2]. As a result, targeted minigene constructs that retain the critical local splicing context often represent a more practical and scalable approach for functional assessment of candidate splice-altering variants. In addition, traditional minigene workflows have largely relied on agarose gel electrophoresis and Sanger sequencing of individual splice products[13], limiting resolution of complex transcript isoforms. The integration of commercial gene synthesis and long-read single-molecule sequencing now enables rapid generation of locus-specific minigene constructs together with full-length characterisation of splice outcomes in a single assay.

### Designing minigene constructs

Most human genes are over 65kb and contain an average of ten introns, averaging 6355 bp each [2] This is outside of the capability of gene fragment synthesis [14], [15], [16] and transfecting large plasmids can be toxic to cells [17]. Therefore, the genomic content included in the minigene plasmid needs to be carefully considered. The design of the genomic insert is a critical determinant of minigene assay performance, as sufficient sequence context must be preserved to allow physiologically relevant splice site recognition and competition.

The important elements to consider for construct design are the canonical 5’ splice site, canonical 3’ splice site, BP and polypyrimidine tract, along with cis-regulatory elements ESE, ESI, ISE and ISI. The 5′ splice site is located at the start of the intron, while the BP, polypyrimidine tract, and 3′ splice site are arranged in this order typically within the final ~40 nucleotides[18], [19] (Figure 1). Distant branch points located up to 400 nucleotides upstream from the 3’ splice site are rare, occurring in approximately 1% of introns[20]. Exonic regulators (enhancers and silencers) are enriched near exon–intron boundaries, typically within 125 nucleotides of exon ends [21], [22], [23], with intronic regulators, similarly, predominantly constrained to the first and last 150-200 bp of the introns [21], [24].

Therefore, to capture the majority of *cis*-splicing elements a minigene gene insert should contain the exons on either side of the putative splicing variant, along with an abridged intron that contains at least 200-300bp of sequence at both intron-exon boundaries[11].

Splicing outcomes are commonly assessed by RT-PCR followed by visualisation of splice isoforms using agarose gel electrophoresis and/or sequencing of splice products. Sanger sequencing has traditionally been used to characterise splice junctions; however, resolution of individual isoforms often requires size fractionation by agarose gel electrophoresis and extraction of discrete bands when multiple products are present. In contrast, single molecule long-read sequencing approaches (e.g. Oxford Nanopore Technologies) enable full-length cDNA transcript sequencing in single reads, allowing the complete splicing context of each transcript to be resolved. Combinations of exon skipping, partial exon usage, and cryptic splice site activation events can be resolved by long-read single molecule sequencing making this method well suited to minigene-based transcript assays.

In summary, we describe a minigene-based strategy for functional validation of splice-altering variants that leverages precise construct design, commercial gene synthesis, and long-read sequencing on a portable sequencer. This approach enables rapid assay generation and characterisation of splice outcomes, providing a robust alternative when patient-derived RNA is unavailable or insufficient.

## Step-by-step protocol

### 1. Design gene fragment

Timing: Variable

The gene fragment should include the candidate splicing variant of interest together with sufficient flanking sequence to preserve cis-acting splicing elements. At a minimum, this should encompass two exons (at least one downstream, 3’ to the variant of interest) and sufficient intervening intronic sequence to enable proper exon definition and splice site recognition (Figure 2). An additional 3’ exon will need to be incorporated into the construct if exon skipping is predicted. The full intervening intron sequence is typically not possible, or necessary[1], [2]. Inclusion of approximately 200-300 bp of intronic sequence at both the 5′ and 3′ ends of each exon is generally sufficient to retain the majority of ISEs and ISSs. For intronic variants, a similar genomic region (200-300 bp on either side of the variant) should be included (unless already captured within the previously mentioned intronic regions adjacent to intron-exon boundaries). A corresponding wild-type construct should also be designed, differing only at the variant position, to enable direct comparison of splicing outcomes.

**Figure 2.**
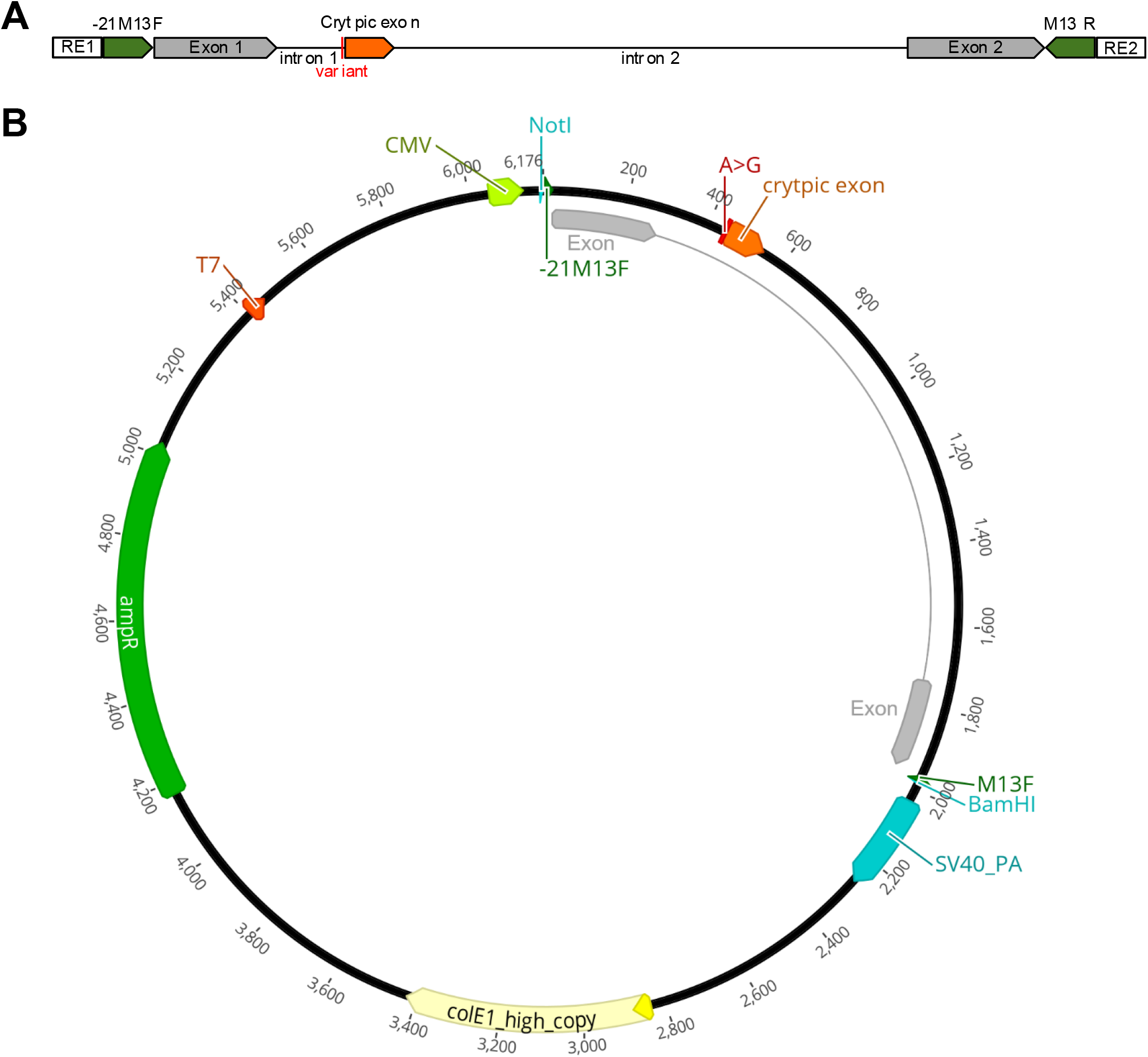
Example minigene construct. A. Example of a gene insert designed and synthesised for validating the effect of a predicted splicing variant *in vitro*. Restriction sites () on the 5’ and 3’ ends are selected to allow for ligation into an expression plasmid. For this construct these are Not1 and BamH1. Primer sites (denoted in dark green) allow for selective amplification of the gene insert for downstream analysis. The variant of interest in this case is a splice acceptor within an intron, which is predicted to result in a cryptic exon. The two bordering exons are also included in the designed gene fragment. B. Gene fragment in the expression plasmid (pTwist CMV) under control of the strong CMV promoter.

Downstream splicing outcomes are assessed by cDNA synthesis from RNA isolated from the plasmid transfected cells, PCR amplification, and subsequent amplicon sequencing. To ensure selective amplification of minigene-derived transcripts expressed in human cells, primers are designed to anneal to vector-derived sequences flanking the insert rather than to the genomic sequence. For example, commonly used vector primers such as −21M13F and M13R can be used where included in the construct design. This approach enables specific amplification of transcripts originating from the minigene while avoiding amplification of transcripts derived from the cellular genome and allows detection of alternative splice products generated from the inserted genomic fragment.

While this protocol recommends outsourcing minigene construct generation to commercial providers, these workflows still rely on standard cloning approaches such as restriction digestion and ligation to insert your minigene sequence into your vector of choice. Accordingly, the gene fragment should be designed with appropriate restriction sites at its termini to facilitate cloning, ensuring that these sites are unique and not present elsewhere within the fragment.

Finally, for most applications, the selection of an appropriate expression vector will be a high-copy number, high-expression vector with a strong promoter such as CMV (e.g. pcDNA3.1(+) or pTwist CMV). Where bacterial amplification of the plasmid vector is required, inclusion of an antibiotic resistance marker is necessary. However, modern gene synthesis services provide constructs as sequence-verified plasmid DNA at scales suitable for downstream applications and at relatively low cost, reducing the need for further cloning or plasmid expansion; as such, antibiotic resistance would not be required [14], [15], [16].

Plasmids can be ordered from a commercial provider at a quantity sufficient for the transfection experiments (e.g. ≥2.5 µg).

Note: Examples of suitable commercial providers include Twist Bioscience, GenScript, and Integrated DNA Technologies (IDT). Choice of provider may depend on cost, turnaround time, and construct complexity, with options such as increased yield or endotoxin-free preparations available. We used pTwist CMV from Twist Bioscience.

### 2. Cell Culture

Timing: ~60-72 hours

Where possible, select a cell line that is representative of the tissue or cell lineage that is affected in disease. From a technical perspective, minigene assays that involve transient transfection perform best in adherent cell lines that exhibit rapid growth and high transfection efficiency, such as HEK-293 cells.

1. Seed 5.0 x 10^5^ of the desired cell line in 2ml of complete DMEM in a 6 well plate
2. Culture for 24 h at 37°C, 5% CO_2_ to reach 70%-90% confluency at transfection
3. Dilute 7.5 µL Lipofectamine 3000 Reagent in 125 µL OptiMEM Medium to make the Transfection Reagent
4. In a separate tube, dilute 2.5 µg plasmid DNA and 5 μL P3000 reagent up to a final volume of 125 μL with OptiMEM
5. Combine 125 μL of Transfection Reagent with the diluted DNA. Incubate together for 5 min at room temperature
6. Add the DNA-lipid complex directly to the media of the cultured cells and swirl to distribute
7. Remove media after 6 hours and replace with fresh complete DMEM
8. Harvest cells 24–36 h after transfection

Note: These conditions that have been successful with a variety of plasmids transfecting HEK-293 cells. While, this protocol suggests the use of lipofection, electroporation is also a possible technique that could be used for plasmid transfection. For optimal transfection and expression, plasmid concentration should be optimised for each plasmid and cell line.

### 3. Extract RNA and Reverse transcribe RNA

Timing: ~4 hours

1. Extract total RNA from transfected cells with a standard RNA extraction kit Note: We use the QIAGEN RNeasy Mini Kit according to the manufacturer’s instruction (https://www.qiagen.com/us/resources/kithandbook/hb-0435-007-hb-rnymini-0623-ww)
2. Synthesize cDNA from RNA using Invitrogen SuperscriptIII First-Strand Synthesis SuperMix kit (https://documents.thermofisher.com/TFS-Assets/LSG/manuals/superscript_firststrandsupermix_man.pdf)

a. Combine the cDNA synthesis reaction components specified in the kit manual:

b. Incubate in a thermal cycler at 65°C for 5 minutes, and then immediately place on ice for at least 1 minute. Collect the contents of the tube by brief centrifugation
c. Add 10 μl 2X First-Strand Reaction Mix and 2 μl SuperScript III/RNaseOUT Enzyme Mix to the cDNA synthesis reaction
d. Vortex the sample briefly to mix and collect by brief centrifugation
e. Incubate at 50°C for 50 min
f. Terminate the reactions at 85°C for 5 minutes. Chill on ice.

| cDNA Synthesis Reaction |  |
| --- | --- |
| Component | Volume |
| RNA | up to 5 µg |
| Primer – 50uM oligo(dT<sub>20</sub>) | 1 ul |
| Annealing buffer | 1 ul |
| Nuclease-free water | to 8 ul |

Note: We use the Invitrogen SuperscriptIII First-Strand Synthesis SuperMix kit according to the manufacturer’s instructions, but other cDNA synthesis kits are available.

### 3. Amplify spliced gene fragment from cDNA

Timing: ~2 hours

1. Amplify minigene-derived transcripts from cDNA by PCR using primers targeting sequences unique to the construct as designed in step 1 (https://www.sigmaaldrich.com/deepweb/assets/sigmaaldrich/product/documents/304/660/2grhsrmkb.pdf).
  a. Combine KAPA2G Robust HotStart ReadyMix PCR Reaction Mix:

  b. Amplify cDNA amplicon using the following thermocycler conditions:

| KAPA2G Robust HotStart ReadyMix PCR Reaction Mix |  |  |
| --- | --- | --- |
| Component | Final concentration | Volume (one sample) |
| Template cDNA | 1-100 ng |  |
| 10 µM -21M13 Forward | 0.5 µM | 1.25 |
| 10 µM M13 Reverse | 0.5 µM | 1.25 |
| 5X KAPA2G Robust HotStart ReadyMix | 1 x | 12.5 |
| PCR-grade H2O |  | To 25 ul |

2. Resolve PCR products by agarose gel electrophoresis to assess splice isoform size and complexity
  a. Load 10 μl of the PCR product mixed with 1 μl of 6x Loading Dye onto a 1.0– 1.5% agarose gel

- b. Amplify cDNA amplicon using the following thermocycler conditions:
| KAPA2G Robust HotStart ReadyMix PCR Thermocycler conditions |  |  |  |
| --- | --- | --- | --- |
| Step | Temperature | Time | Cycles |
| Initial denaturation | 95°C | 15s | 1 |
| Denaturation | 95°C | 15s | 35 |
| Annealing | 60°C | 15s |  |
| Extension | 72°C | 30s |  |
| Final Extension | 72°C | 1min | 1 |

Note: We use Roche Kappa Robust 2G Readymix with the -21M13F and M13R primers, but primers will be specific to the sequences designed in step 1, and many other DNA polymerase systems are available.

Distinct bands corresponding to alternative splice products may be visualised directly from an electrophoresis gel and compared between the wild type and variant constructs.

### 6. Purify PCR products and sequence

Timing: ~8 hours

Resolving the PCR-derived bands from step 5 on an agarose gel will demonstrate a change in splicing resulting from the variant introduced into the gene fragment. However, exact alterations to the mRNA transcript resulting from the splice variant can be determined using sequencing. We use Oxford Nanopore Technologies (ONT) sequencing to capture the entire strand of amplified cDNA on a single sequence read.

1. Purify PCR products using Beckman Coulter AmpureXP beads (https://www.beckman.com/techdocs/B37419AB/wsr-150180)
  a. Add 0.5x vol (10 ul) of Ampure XP beads to remaining PCR products (20 ul)
  b. Using magnetic rack, wash twice in 80% ethanol
  c. Remove supernatant
  d. Dry beads on magnetic rack for 30s
  e. Remove from rack
  f. Resuspended PCR amplicons in 10 ul PCR grade water for 2 min at RT
  g. Place on magnetic rack. Remove supernatant into sterile microfuge tube
  h. Quantify purified PCR product using Invitrogen Qubit 1X dsDNA Broad Range (BR) Assay (https://documents.thermofisher.com/TFS-Assets/LSG/manuals/MAN0019617_Qubit_1X_dsDNA_BR_Assay_UG.pdf)
2. Sequence amplified transcripts on a flongle flowcell using ONT Native Barcoding Kit 24 V14 (SQK-NBD114.24) according to the manufacturer’s instructions (https://nanoporetech.com/document/ligation-sequencing-amplicons-native-barcoding-v14-sqk-nbd114-24)

Note: We used ONT Native Barcoding Kit 24 V14 (SQK-NBD114.24) which contains 24 barcodes. If you have more amplicons and need more barcodes, Native Barcoding Kit 96 V14 (SQK-NBD114.96) is available with up to 96 barcodes.

Note: barcoding via PCR is also possible for ONT using the method described by Li, et al. (2024) [25].

## 6. Align and analyse sequence

Timing: 1 hour

1. Generate fasta reference for the gene fragment
  a. An example of a “chromosome” for the reference in Figure 3.
2. Perform basecalling and inline barcode separation pod5 files from ONT sequencing using Dorado duplex basecalling (https://software-docs.nanoporetech.com/dorado/latest/basecaller/duplex/)
  a. Use the command `dorado duplex hac pod5s/ --kit-name SQK-NBD114.24 > calls.duplex.bam`
3. Align the barcoded sequences to the reference using Dorado (https://software-docs.nanoporetech.com/dorado/latest/basecaller/alignment/)
  a. Use the command `dorado aligner index.fasta calls.duplex.bam > aligned.bam`
4. View alignments using Integrative Genomics Viewer (IGV, https://igv.org/) (Figure 4A)
  a. Generate a Sashimi plot to quantify differences in splicing resulting from splicing variants (Figure 4C)

**Figure 3.**
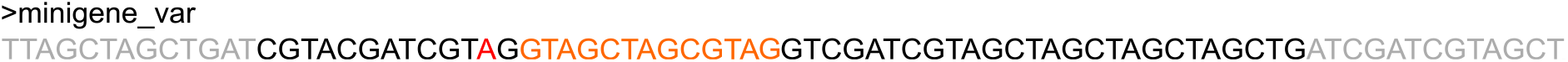
Example fasta reference for read alignment. Your reference file will be a plain text file, however, we have coloured specific parts of a condensed fasta to demonstrate the parts of the gene fragment that should be included in the reference file. Grey represents the exons and black the introns. Red represents the predicted splice acceptor variant used as an example here, with orange representing the predicted cryptic exon that will result from altered splicing.

**Figure 4.**
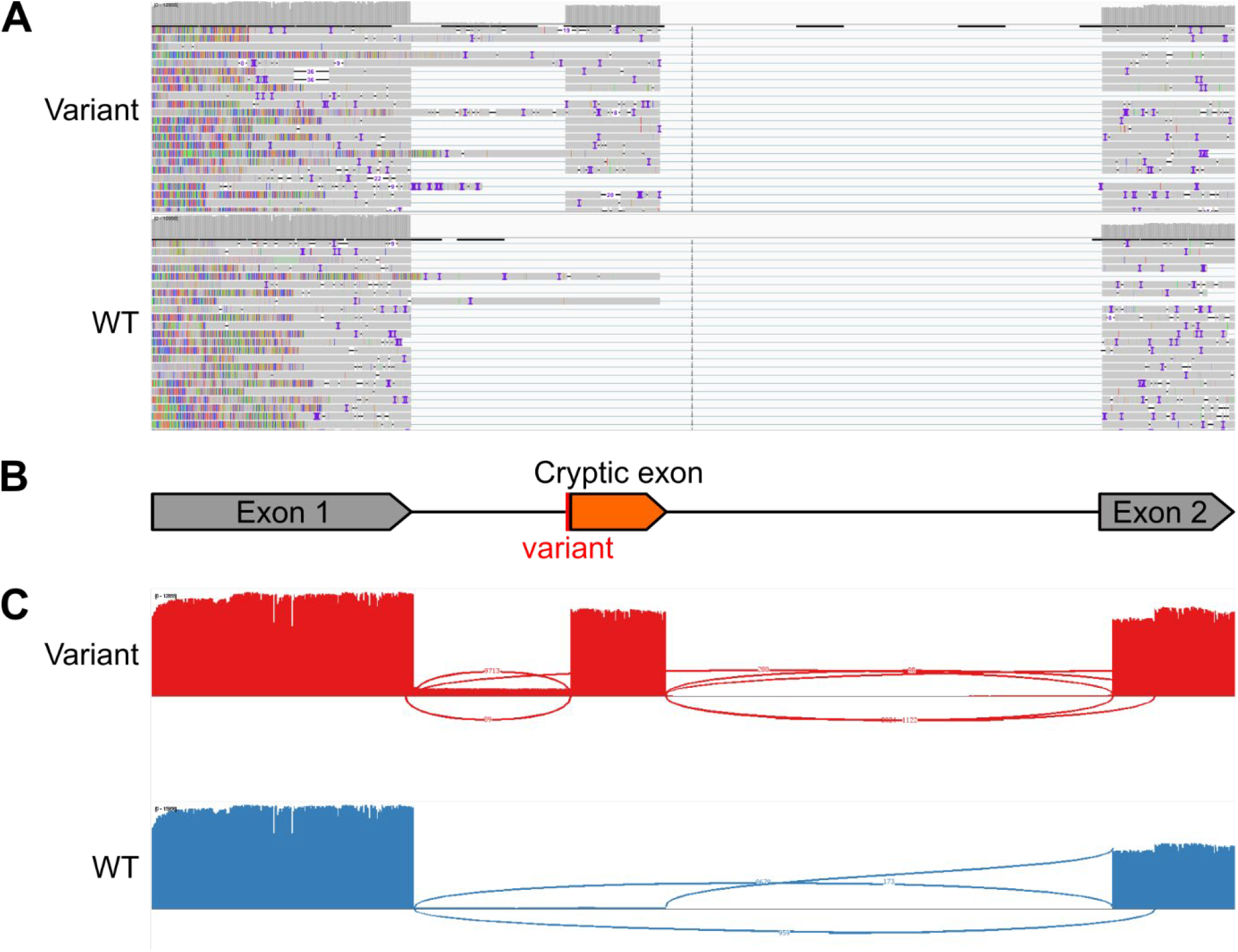
Example of aligned ONT sequencing results of alternate splicing exons. A. Coverage track and read alignments for ONT PCR amplicon sequencing of cDNA of the minigene gene fragment after expression in cultured cells. Coverage track demonstrates the inclusion of the predicted cryptic exon in the mature mRNA resulting from the putative splice acceptor variant introduced in the variant gene fragment only, and not in the sequence from the wild type (WT) gene fragment. B. Genomic component of the gene fragment which represents the sequence in the reference file (Figure 3). The grey arrow represents the exons included in the fragment, with the black line representing the introns. The candidate splice variant is indicated by the vertical red line, with the predicted cryptic exon indicated by the orange arrow. C. Sashimi plot of splicing in the ONT PCR amplicon sequencing of cDNA of the minigene gene fragment after expression in cultured cells. Consistent with the coverage and alignment tracks, the Sashimi plot shows that the majority of transcripts resulting from the variant gene fragment displays altered splicing, with the mature mRNA including exon 1, the cryptic exon and exon 2 spliced together. In comparison, the transcripts from the WT gene fragment result in mature mRNA that has exon 1 and exon 2 spliced together without the cryptic exon.

## Expected outcomes

Successful execution of this protocol should result in robust expression of minigene-derived transcripts that can be specifically amplified from host cell RNA. cDNA PCR amplification typically produces one or more discrete bands corresponding to alternative splice isoforms. For wild-type constructs, a dominant band representing canonical splicing is expected, although minor alternative isoforms may be observed depending on gene context.

Variant constructs may show altered splicing patterns, including exon skipping, partial exon inclusion, intron retention, or activation of cryptic splice sites. These changes may manifest as shifts in band size on agarose gel electrophoresis or as additional bands relative to the wild-type construct.

Long-read sequencing enables full-length characterisation of individual transcript isoforms in single reads, allowing unambiguous resolution of splice junctions and combinations of splicing events (Figure 5). This is particularly important where multiple isoforms are present or where complex splicing patterns arise. Quantitative differences in isoform abundance from the same variant may also be assessed, important in cases where the genetic variant results in alternative splicing where multiple mature mRNAs result from the same nascent transcript.

**Figure 5.**
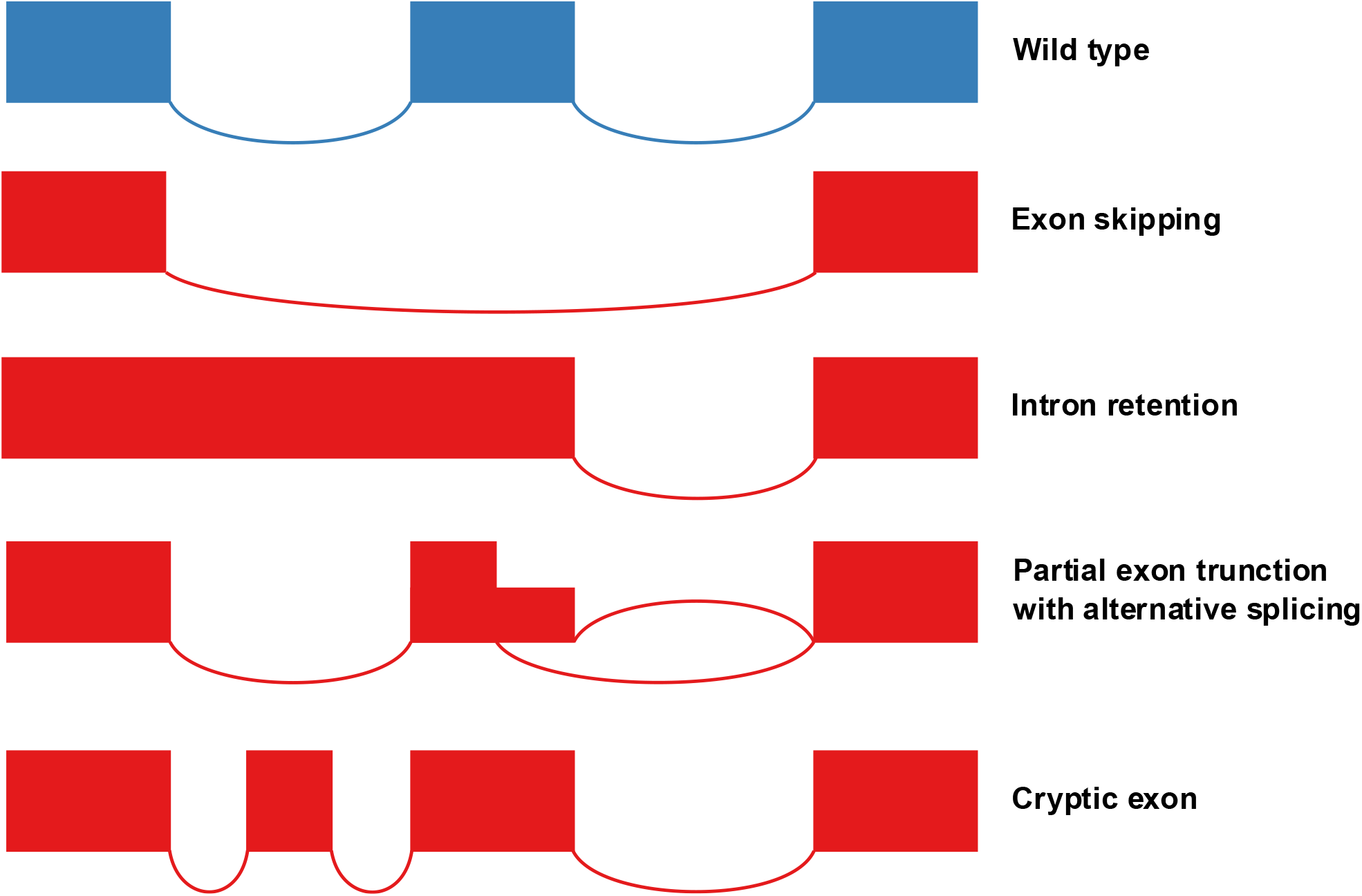
Representative splicing outcomes of splice-altering variants. Example Sashimi plots illustrating common patterns of aberrant splicing, including exon skipping, intron retention, partial exon truncation with alternate splicing, and activation of cryptic splice sites.

Overall, comparison between wild-type and variant constructs enables direct assessment of the functional impact of the variant on splicing.

## Limitations

Minigene assays provide a controlled and reproducible system for assessing splicing, however, they do not fully recapitulate endogenous genomic context. Regulatory elements located outside the cloned region, including distant intronic or chromatin-associated features, are not captured and may influence splicing *in vivo*.

Splicing outcomes may also be influenced by the cellular context. The use of heterologous cell lines (e.g. HEK-293) may not reflect tissue-specific splicing patterns, particularly for genes with tissue-specific expression or regulation.

Insert design represents a critical limitation. Although inclusion of ~200–300 bp of flanking intronic sequence captures the majority of proximal regulatory elements, more distal elements may be missed, potentially leading to incomplete or misleading splicing outcomes.

## Troubleshooting

### Problem 1: No or low PCR product

#### Possible causes

Low transfection efficiency; poor RNA quality; low expression of the construct; inefficient primer design.

#### Potential solutions

- Confirm transfection efficiency (e.g. using a control plasmid)
- Verify RNA integrity and concentration
- Confirm cDNA synthesis using control RNA (often included in cDNA synthesis kits)
- Increase input RNA or cDNA
- Optimise PCR conditions (annealing temperature, cycle number)
- Redesign primers targeting construct-specific sequences

### Problem 2: Amplification of non-specific or endogenous transcripts

If unexpected bands are present in the wild type PCR product, this indicates amplification of regions that are non-specific to the gene fragment.

#### Possible causes

Primers annealing to endogenous sequences; insufficient specificity of primer design.

#### Potential solutions

- Ensure primers target sequences unique to the construct
- Optimise PCR conditions (e.g. alter annealing temperature with a temperature gradient)

### Problem 3: Multiple unexpected bands

#### Possible causes

Alternative splicing events; incomplete construct design; PCR artefacts.

#### Potential solutions

- Confirm isoforms in sequencing data
- Reduce PCR cycle number to minimise artefacts

## Resource Availability

### Lead contact

Further information and requests for resources should be directed to the lead contact, Whitney Whitford.

### Technical contact

Questions about the technical specifics of performing the protocol should be directed to technical contact, Whitney Whitford.

### Materials availability

No new unique reagents were generated in this study. However, further requests for resources should be directed to the lead contact, Whitney Whitford.

### Data and code availability

This study did not generate code, and no specific sequence was generated as protocol serves to discuss general methodology.

## Acknowledgments

W.W. was supported by the Neurological Foundation of New Zealand through the Dawn Fellowship.

## Author contributions

W.W. and S.M. performed experiments to establish the protocol. W.W. conceived the project and W.W. J.C.J, and R.G.S. were involved in methodology refinement. W.W. wrote the first draft. All authors participated in review and editing. All authors have read and agreed to the content.

## Key Resources Table

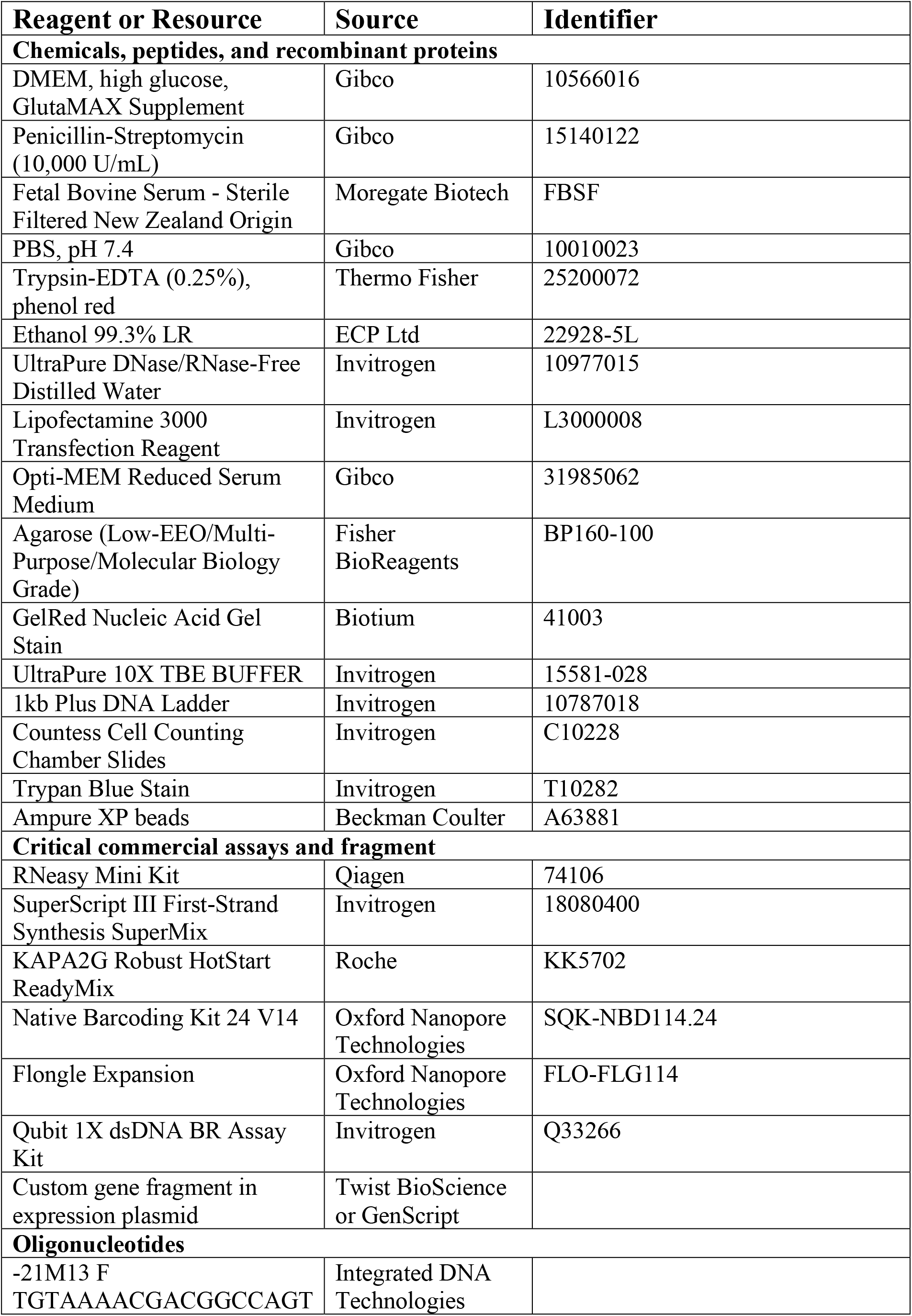
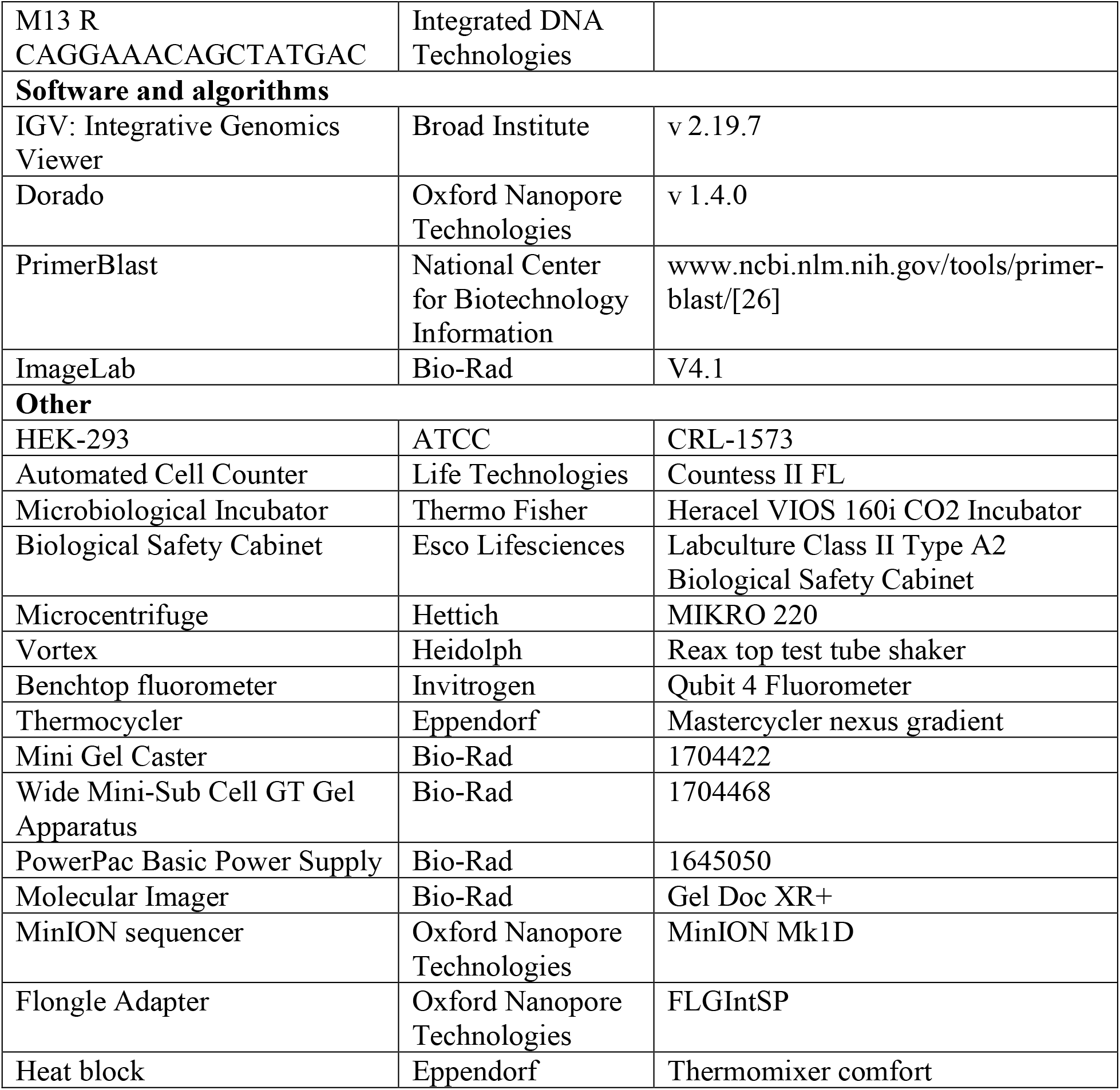

## Materials and Equipment

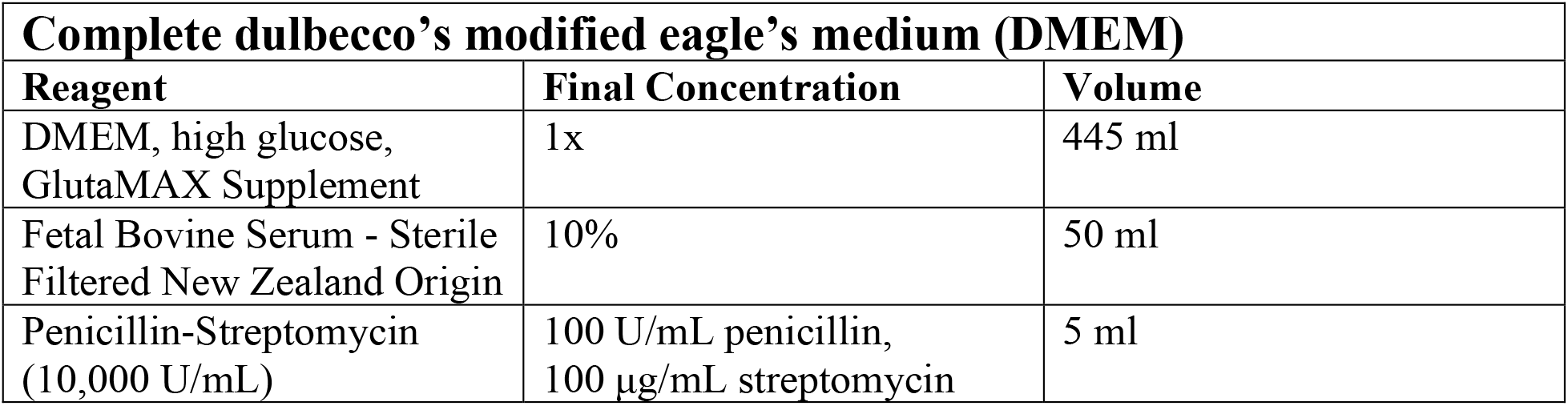

| **Complete dulbecco's modified eagle's medium (DMEM)** |  |  |
| --- | --- | --- |
| **Reagent** | **Final Concentration** | **Volume** |
| DMEM, high glucose,<br>GlutaMAX Supplement | 1x | 445 ml |
| Fetal Bovine Serum - Sterile<br>Filtered New Zealand Origin | 10% | 50 ml |
| Penicillin-Streptomycin<br>(10,000 U/mL) | 100 U/mL penicillin,<br>100 µg/mL streptomycin | 5 ml |

